# The bacterial community of the European spruce bark beetle in space and time

**DOI:** 10.1101/2023.04.28.538755

**Authors:** Abdelhameed Moussa, Stefano Nones, Patrizia Elena Vannucchi, Gul-i-Rayna Shahzad, Jessica Dittmer, Erika Corretto, Martin Schebeck, Massimo Faccoli, Andrea Battisti, Christian Stauffer, Hannes Schuler

**Author notes:** Corresponding author: Hannes Schuler, Competence Centre for Plant Health, Free University of Bozen-Bolzano, Bozen-Bolzano, Italy.

## Abstract

The European spruce bark beetle *Ips typographus* is a pest causing severe damages in forests dominated by the Norway spruce in Europe. Microorganisms play an essential role in the host species performance, including nutrition, fitness as well as in overcoming host defenses. Here, we performed high-throughput 16S rRNA metabarcoding of *I. typographus* across different populations in Europe, to assess its bacterial community. We investigated four postglacial refugial areas in Europe and focused specifically on a current bark beetle hot spot in the Dolomites where we compared populations with different epidemiological phases (outbreaking vs. non-outbreaking) and across different seasons (pre-overwintering vs. overwintering). Our results show that the bacterial community structure varied among populations from the refugial areas and between different regions within the Dolomites. We found a significant difference in the bacterial community between pre-overwintering and overwintering individuals, but we did not find differences between epidemic and endemic populations. The prevalence of the genus *Erwinia* which was present in every individual and *Pseudoxanthomonas* in almost every individual across all populations, suggests that these taxa form the core bacterial community of *I. typographus*. Furthermore, several additional bacterial taxa occurred in all populations, but with variable frequencies. This study highlights a complex interaction of *I. typographus* and various bacterial taxa across different regions and ecological phases of *I. typographus* populations and provides new insights into the role of microorganisms in the biology of this important pest species.

## Introduction

The European spruce bark beetle *Ips typographus* (L.) (Coleoptera: Curculionidae, Scolytinae) is one of the most destructive forest pests in Europe, which has caused significant ecological and economic disturbances over the last decades (Schelhaas et al. 2003). In Europe, *I. typographus* has a wide geographic range overlapping with its main host tree species, Norway spruce *Picea abies* (Karsten). Normally *I. typographus* exist as endemic populations were beetles breed and develop within the phloem of weakened and dying trees, especially wind-thrown and drought-stressed trees with impaired defenses. In epidemic phases when population outbreaks occur, this beetle can colonize and kill also apparently healthy trees (Schroeder 2010). In 2018, the storm Vaia destroyed more than 410 km^2^ of forests, mainly dominated by Norway spruce, in the Southern Alps of North-Eastern Italy and Southern Austria (Udali et al. 2021). Moreover, heavy snow falls affected the Dolomites region in the years 2019 and 2020. These forest disturbance events provided vast amounts of suitable host material for *I. typographus* reproduction and development and resulted in a dramatic bark beetle outbreak in the Southern Alps (Nardi et al. 2022).

Insects are associated with various microorganisms that provide physiological and ecological benefits to their hosts. They play a significant role in various aspects of insect life histories, including nutrition, development, morphogenesis, behavior, and immunity (Douglas 2015). In particular, microorganisms are of high significance for phytophagous insects. For instance, despite having endogenous detoxification systems, insects are often associated with microbes to detoxify protective plant secondary metabolites (Itoh et al. 2018).

Bacterial symbionts also play a significant role in the life histories of numerous bark beetles, such as in the economically important genera *Ips* and *Dendroctonus*, by contributing to the detoxification of plant metabolites, protection of beneficial fungal symbionts, nutrition, detoxification, production of pheromone-like molecules, and protection against pathogens through the production of antimicrobial compounds (García-Fraile 2018). For instance, the bacterial symbiont *Erwinia typographi* associated with *I. typographus* showed resistance to high concentrations of the spruce monoterpene myrcene, highlighting a potentially important role of this bacterium in the defense against terpenoids and phenolic compounds and thus assisting in overcoming host defense mechanisms (Skrodenytė-Arbačiauskienė et al. 2012). The genus *Pseudomonas* has been isolated from different *I. typographus* ontogenic stages and was shown to play a potential role in nutrient provisioning, protection against pathogens, and degradation of toxic compounds (Peral-Aranega et al. 2020). Therefore, bacteria might assist in the successful colonization of trees and might therefore play an important role in mass outbreakings of *I. typographus*.

Although our understanding of the bacterial community structure of *I. typographus* has increased in recent years (Chakraborty et al. 2020; 2023; Fang et al. 2020; Yu et al. 2022; Veselská et al. 2023), surprisingly little knowledge exists about the taxonomic composition and diversity across different geographic regions, epidemiological and overwintering phases. Here, we first describe the taxonomic composition and core-microbiome of *I. typographus* by characterizing the bacterial communities of different populations from a large area of Europe, including the four major refugial areas, in the Apennines, the Dinaric Alps on the Balkan Peninsula, the Carpathian Mountains and in the Russian plain (Tollefsrud et al. 2009). Subsequently, we focus particularly on the Dolomites (Northeastern Italy and Southern Austria), where currently an outbreak of this beetle occurs. In this region, we compare the microbial composition of endemic and epidemic populations to investigate the role of bacteria in the population dynamics of *I. typographus* and investigate the potential role of bacteria in overwintering individuals by comparing the bacterial communities of populations in different overwintering phases.

## Materials and methods

### Sample collection

Adults of *Ips typographus* were collected from seven geographic regions. Populations from the Apennines (Abetone – Italy, Abe), the Dinaric Alps in Croatia (Vrhovine, CR), the Carpathian Mountains in Romania (Belis, RO), and the Russian plain (Alexandrov – Russia, RU) were included to assess differences in the bacterial community across a broad geographic scale and evaluate potential effects of historic events during the Pleistocene (Figure 1; Table S1). Moreover, we focused specifically on populations within the Dolomites that were affected by the storm Vaia in 2018 and by the snowfall events in the following years, namely four populations from Eastern Tyrol (Austria; ET1-ET4), three populations from South Tyrol (Italy; ST1-ST3) and six populations from Veneto (Italy; VT1-VT6) (Figure 1). To compare the bacterial communities between epidemic and endemic populations on a small local scale, the four populations in Eastern Tyrol were collected in different epidemiological phases in summer 2020 (two endemic sites, ET2 and ET4, and two epidemic sites, ET1 and ET3). The populations from Veneto were sampled before and during winter of 2019 and 2020, to compare the microbial community from pre-overwintering and overwintering individuals. For each population, 8-12 individuals were collected from infested *Picea abies* trees (Suppl table S1). To avoid the analysis of siblings, single adult beetles were collected from different galleries. Live beetles were immediately transferred to absolute ethanol and stored at -20 °C. Detailed information about the locations is listed in Table S1.

**Figure 1:**
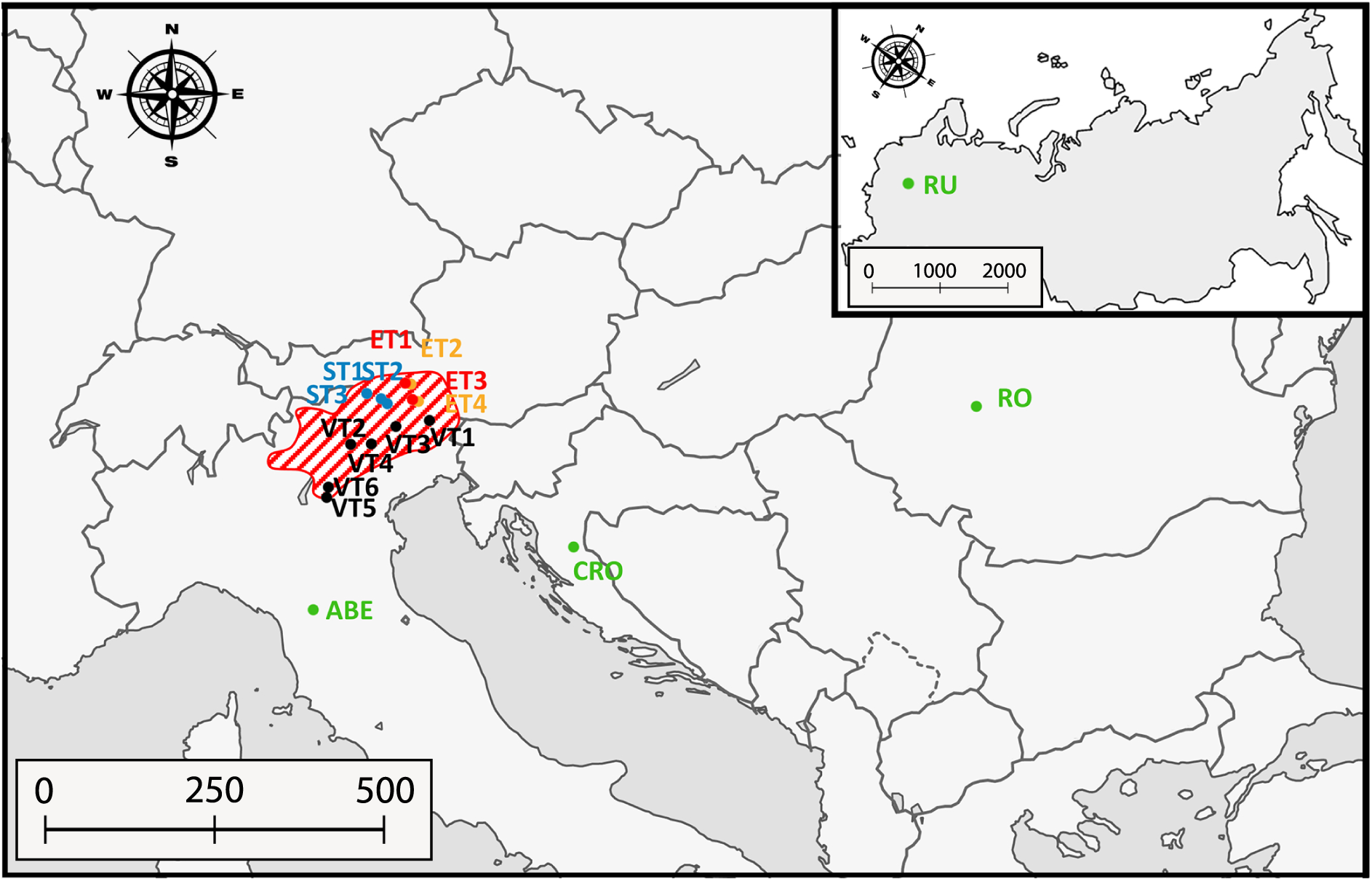
Overview of *Ips typographus* sample locations. ET1-4: Eastern Tyrol; ST1-3: South Tyrol; VT1-6: Veneto; ABE: Abetone; CR: Croatia; RO: Romania and RU: Russia. Populations depicted in orange represent populations in endemic phase, whereas red indicates epidemic phase. Populations in green represent populations sampled before overwintering, whereas populations in blue were sampled during overwintering. The dashed background shows the area affected by the current *I. typographus* outbreak in the Southern Alps. Details of population localities are given in Table S1.

### DNA extraction and sequencing

DNA was extracted from single *I. typographus* individuals using the Sigma-Aldrich GenElute Mammalian Genomic DNA Miniprep Kit (Saint Louis, Missouri, USA) following the manufacturer’s protocol. DNA was quantified with the Invitrogen™ Qubit® dsDNA High Sensitivity (HS) Assay Kit (Life Technologies, USA) and DNA quality was checked with a Nanodrop 2000 (Thermo Scientific, Waltham, Massachusetts, USA). As our aim was to investigate the whole bacterial community associated with *I. typographus*, the collected individuals were not surface-sterilized prior DNA extraction. The bacterial communities of 192 individuals were amplified using barcoded primers 515f and 806r which amplify the V4 region of the 16S rRNA gene and sequenced on an Illumina MiSeq platform using 2×250 bp paired-end chemistry by a commercial provider (StarSEQ GmbH, Mainz, Germany). Moreover, four no template controls containing ultrapure H_2_O were included as negative controls and a mock community including ten bacterial taxa was included as positive control. Raw reads were deposited at NCBI with the BioProject ID PRJNA1020540.

### Data analysis

Raw reads were analyzed using the QIIME2 pipeline (version 2021.4). Due to the poor quality of the reverse reads, only the forward reads (which span almost the entire V4 amplicon) were used for the subsequent analyses. DADA2 was used for quality filtering, denoising, and Amplicon Sequence Variant (ASV) calling using the plug-in q2-dada2 with the denoise-single method. Taxonomic assignment was done against the SILVA database version 138 (Quast et al. 2012). ASVs identified as chloroplasts, mitochondria, archaea, and eukaryotes were filtered out along with rare ASVs (i.e., singletons and ASVs represented by < 10 reads). Moreover, all ASVs present in the negative controls were removed, besides one ASV corresponding to *Erwinia* which was present in low abundance in one of the four controls, but in high abundance across all individuals. The resulting ASVs table (Table S2) was then used for the subsequent analyses.

Data analysis was done using R (version 4.1.2) using the ‘phyloseq’ package (McMurdie & Holmes 2013). Shared and unique ASVs of the different geographic locations were identified, considering only ASVs with 10 or more reads per sample. Visualization of the intersecting and unique ASVs was carried out using the ‘UpSetR’ package (Conway et al. 2017).

Bacterial diversity was analyzed using the packages ‘phyloseq’ (McMurdie & Holmes 2013) and ‘vegan’ (Oksanen et al. 2015), after data normalization using the relative log expression (RLE) method (Robenson & Oshlack 2010). The alpha diversity indices Chao 1 and Shannon’s diversity index as well as Good’s coverage (to validate the representativeness of the sequencing effort) were determined for each sample. Normal distribution of data was evaluated using the Shapiro-Wilk test. ANOVA and Tukey’s Honest Significant Difference (HSD) post-hoc test were used to test for differences between bacterial diversity among populations and localities. Additionally, Welch’s t-test was used to test for differences in diversity between endemic and epidemic populations as well as between pre-overwintering and overwintering populations.

Beta diversity was analyzed using Canonical Analysis of Principal coordinates based on Bray-Curtis and unweighted UniFrac distance metrics using the ‘CAPSCALE’ package (Oksanen et al. 2015) to determine differences in community structure among the different geographical locations. Permutation multivariate ANOVA (Adonis) was used to assess differences in community structure among different geographical locations with 1,000 permutations followed by a pairwise comparison. Furthermore, a circular pattern hierarchical clustering of samples collected from the Dolomite region was performed based on both unweighted UniFrac distance metrics and Bray-Curtis dissimilarity using the ‘ggtree’ package (Yu et al. 2017).

Differentially abundant taxa between pre-overwintering and overwintering populations of Veneto were identified using the linear discriminant analysis effect size analysis (LEfSe; Segata et al. 2011) with default settings using the diff_analysis function in MicrobiotaProcess R package.

Metagenome functions were predicted based on the ASVs abundances of the microbiomes using the original python implementation of PICRUSt2 v2.5 (https://github.com/picrust/picrust2) and MetaCyc (Caspi et al. 2010). Results were visualized in the Marker Data Profiling module of MicrobiomeAnalyst version 2.0 (Lu et al., 2023) with default parameters (low count filter: 4 counts, 20% prevalence; low variance filter: 10% based on inter-quartile range; centered log ratio transformation). Differences between populations, regions, seasons, and epidemiological phases were tested for using t-test or one-way ANOVA (Benjamini–Hochberg method, FDR q < 0.05).

## Results

### *Predominant taxa in the bacterial community of* Ips typographus

After quality control and sequence filtering, a total of 6,797,490 reads ranging from 3,738 to 65,356 reads per individual were retained (Table S2). These were clustered into 11–390 ASVs per individual (mean = 124.24 ± 5.34). These ASVs comprised 31 bacterial phyla, 70 classes, 174 orders, 299 families, and 613 genera. The Good’s coverage was greater than 99.5% in all 192 samples and all rarefaction curves reached a plateau, indicating that the sequencing coverage was sufficient to capture the bacterial communities of all samples (Figure S1).

The bacterial community composition across all *I. typographus* individuals revealed ten major bacterial phyla. The most abundant phyla were Pseudomonadota (formerly Proteobacteria, 25.1%), Bacteroidota (formerly Bacteroidetes, 17.93%), Bacillota (formerly Firmicutes, 11.2%), and Actinobacteria (6.18%), while Myxococcota, Bdellovibrionota, Planctomycetota and Abditibacteriota were present at lower abundance across all regions (Figure 2A). At the genus level, 15 bacterial taxa were present in most of the studied regions, including *Acinetobacter, Allorhizobium-Neorhizobium, Brevundimonas, Chryseobacterium, Erwinia, Izhakiella, Mycobacterium, Novosphingobium, Pseudomonas, Pseudoxanthomonas, Sphingobacterium, Spiroplasma, Williamsia* and *Wolbachia* (Figure 2B). The genus *Erwinia* was the most represented genus, being present in all individuals. Moreover, the genus *Pseudoxanthomonas* was present in all but three individuals in Russia (98.44% of all individuals) and occurred in all studied regions (Figure S2). Therefore, *Erwinia* and *Pseudoxanthomonas* can be considered as the core bacteria of *I. typographus*.

**Figure 2:**
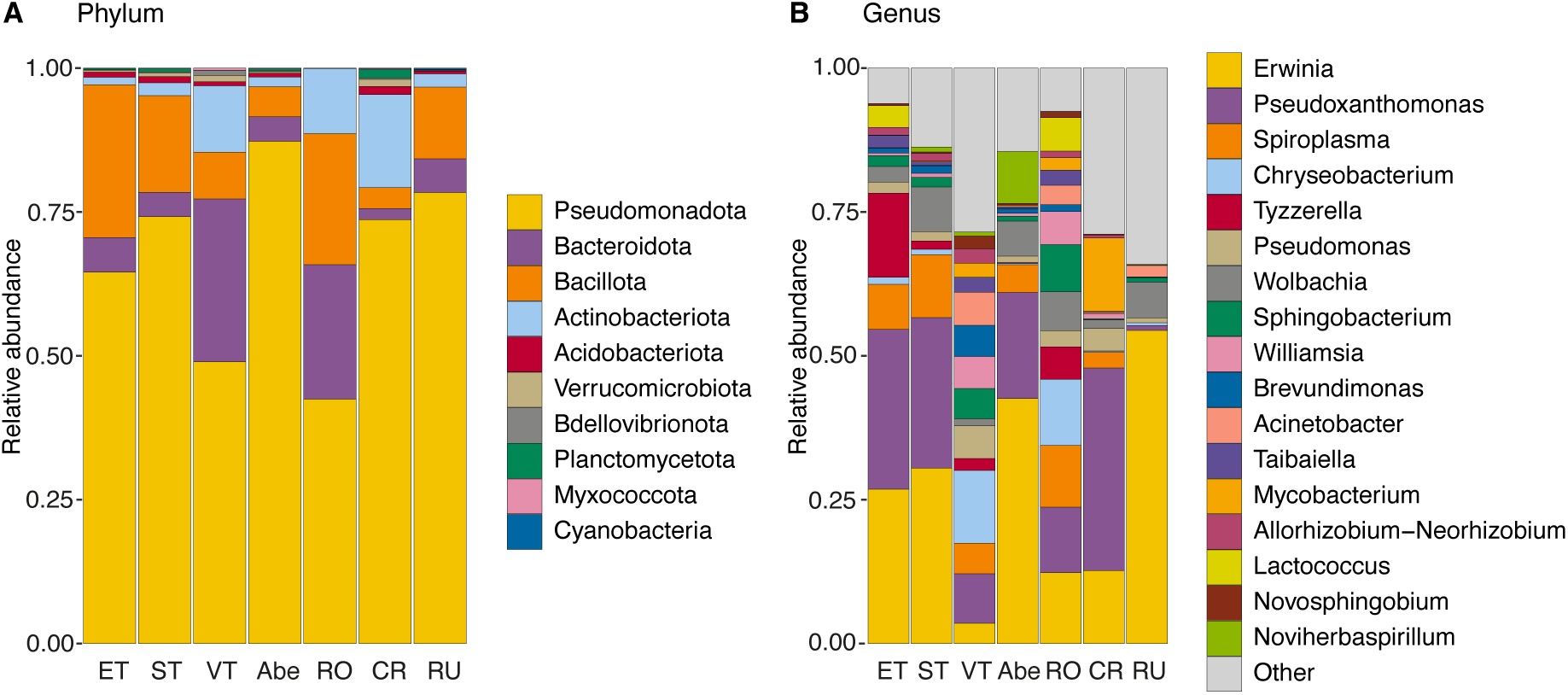
Microbial composition of *Ips typographus* across different populations at phylum level (A) and genus level (B). In (B) only the 17 most abundant genera were plotted.

### Differences in bacterial communities across geographic regions

Overall, 15 ASVs were shared among all populations, which belonged to the genera *Erwinia, Pseudoxanthomonas, Acinetobacter, Spiroplasma, Pseudomonas, Chryseobacterium, Sphingobacterium,* and *Allorhizobium-Neorhizobium*. Additionally, eleven ASVs were common in all populations except Russia (Figure 3) which included the genera *Tyzzerella, Taibaiella, Gordonia, Patulibacter* and *Endobacter.* In contrast, the majority of ASVs were exclusively present only in single regions: 312 in the four populations from Eastern Tyrol, 535 in the three populations from South Tyrol, 763 in the six populations from Veneto, 227 in Abetone, 93 in Romania and 335 in Croatia and 477 in Russia. Most ASVs were exclusively present in single populations (Figure 3). At the genus level, 25 genera were shared among all regions and populations, which included *Acinetobacter, Brevundimonas, Chryseobacterium, Erwinia, Novosphingobium, Prevotella, Pseudoxanthomonas, Pseudomonas, Seratia, Sphingobacterium, Sphingobium, Spiroplasma* and *Williamsia.* Most genera were exclusively present only in single populations: 13 in Eastern Tyrol, 27 in South Tyrol, 26 in Veneto, seven in Abetone, two in Romania, ten in Croatia and 44 in Russia (Figure S3).

**Figure 3:**
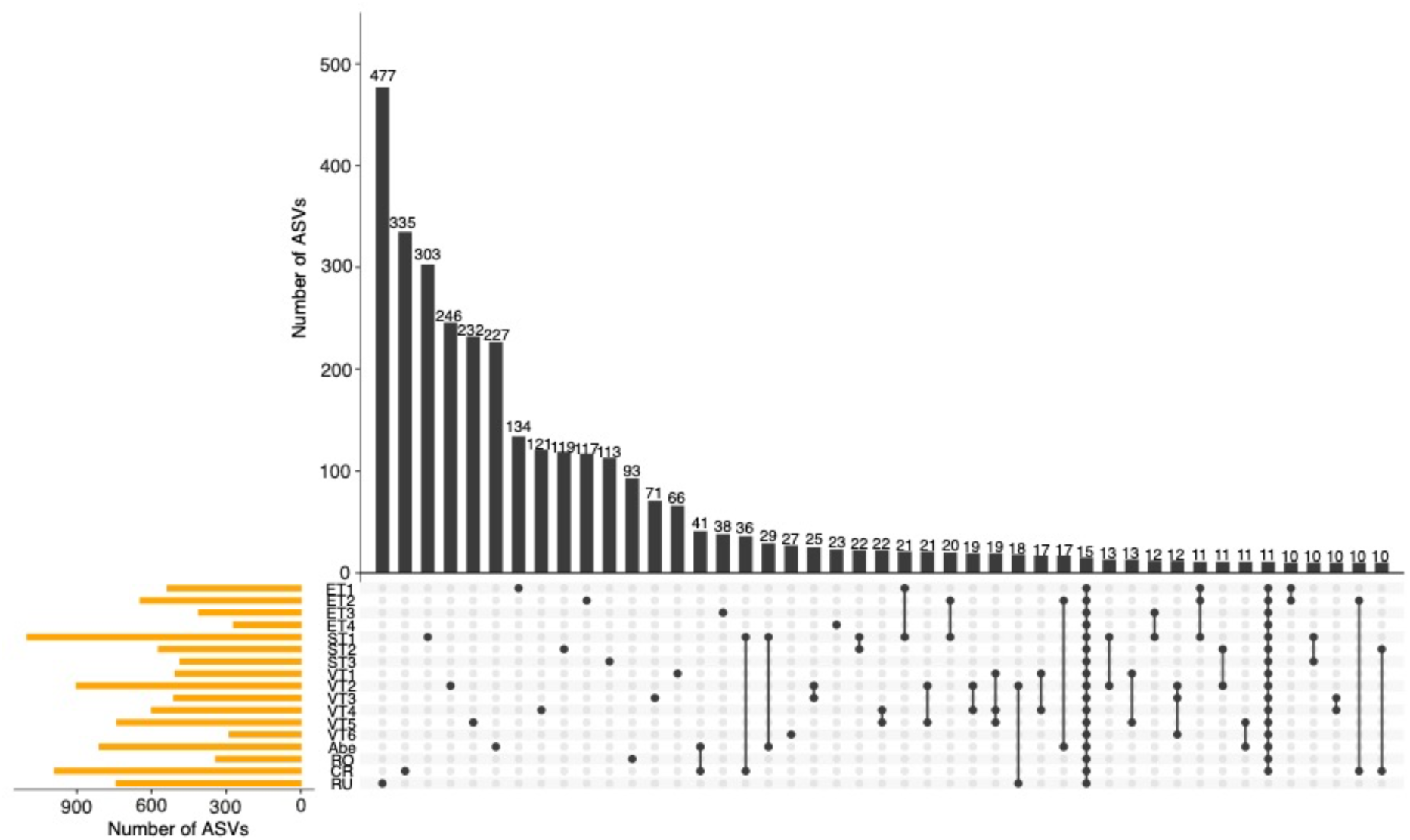
Distribution of ASVs across different populations of *Ips typographus*. Vertical bars (black) and the dot matrix represent the number of shared or unique ASVs, whereas horizontal bars (yellow) show the total number of ASVs present in each population. Only the 40 most common intersections are shown.

Overall, bacterial species richness differed significantly between regions (ANOVA, F= 13.39, df= 6, p < 0.001; Figure 4). Pairwise comparison of species richness showed significant difference between several regions (Table S3). Bacterial diversity based on the Shannon index also differed significantly between regions (ANOVA, F= 20.92, df= 6, p < 0.001; Table S4). Beetles from Veneto had the highest Shannon index of 3.62 whereas individuals from Eastern Tyrol had the lowest of 1.80 (Figure 4). The bacterial community composition of *I. typographus* differed significantly between and within regions. This was confirmed by a distinct segregation and a significant difference between the geographical regions based on unweighted UniFrac distance (PERMANOVA, df= 6, F= 12.733, R^2^= 0.29227, p < 0.001; Figure 5; Table S5). The bacterial species richness was also significant different between the different refugial areas (Chao1: ANOVA, F= 15.07, df= 3, p < 0.0003), whereas bacterial diversity was not significantly different among them (Shannon: ANOVA, F= 1.06, df= 3, p < 0.377).

**Figure 4:**
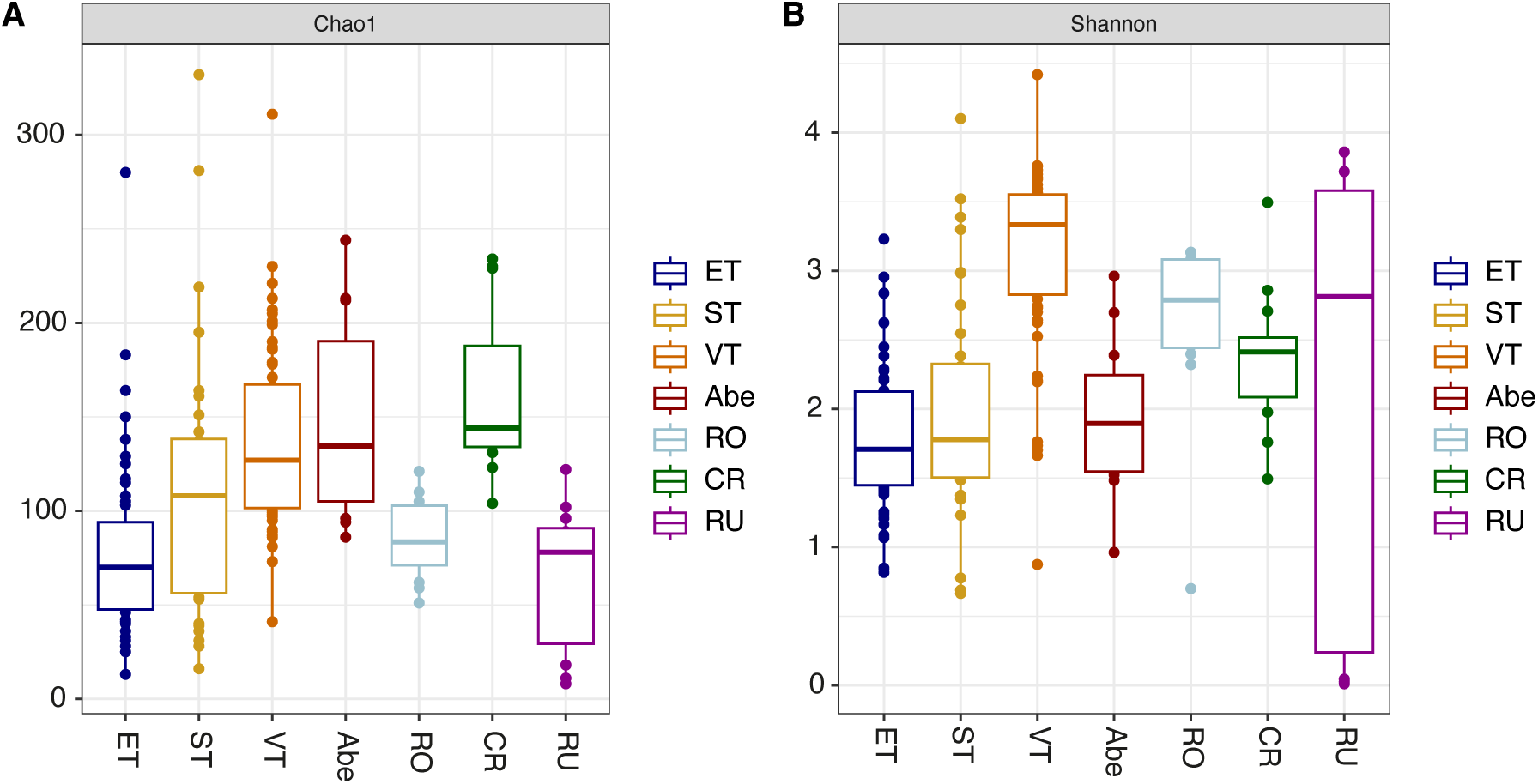
Alpha diversity of the bacterial communities of *Ips typographus* from different geographical regions. Comparison of bacterial species richness (Chao1, A) and bacterial diversity (Shannon index, B).

**Figure 5:**
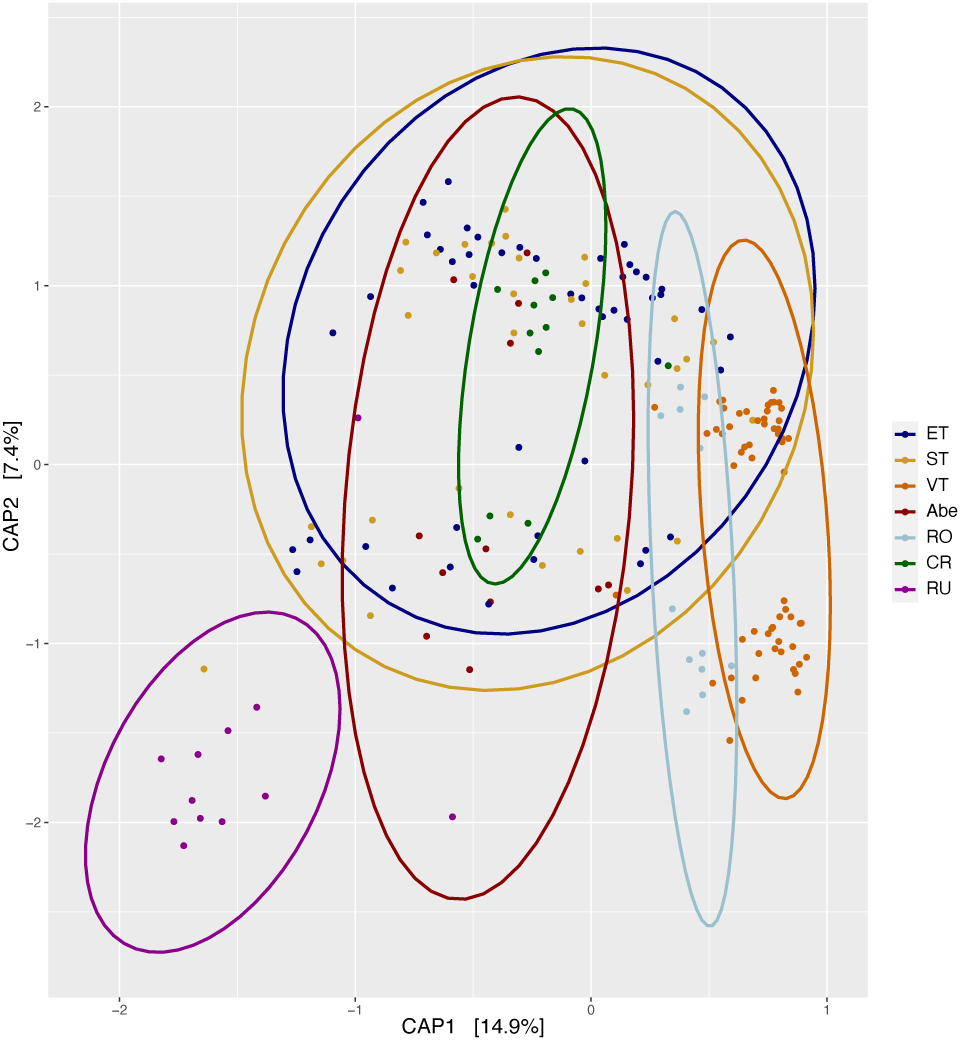
Canonical analysis of principal coordinates (CAP) based on unweighted UniFrac distance showing differences in bacterial community structure of *Ips typographus* among different geographic regions.

The specific focus on populations within the Dolomites allowed the comparison of the bacterial communities of different *I. typographus* populations on a smaller geographic scale. Overall, the bacterial species richness was significantly different between locations from the Dolomites (Chao1: ANOVA, F= 17.2, df= 2, p < 0.001). Pairwise comparisons showed a significant difference between ET and ST (p < 0.02), ST and VT (p < 0.03), and populations from ET and VT had a significantly different bacterial diversity (p < 0.0001; Figure 4). Likewise, bacterial species diversity was significantly different among the Dolomites regions (Shannon: ANOVA, F= 64.28, df= 2, p < 0.0001). Pairwise comparisons showed no significant difference between ET and ST (p = 0.477) but a significant difference between ET and VT (p < 0.0001) and VT and ST (p < 0.0001). These results were confirmed by a PERMANOVA based on unweighted UniFrac distance metrics, which revealed a significant difference in the bacterial community structure between different Dolomite regions (PERMANOVA, df= 2, F= 16.034, R^2^= 0.18529, p < 0.001; Figure 5). However, the pairwise comparison showed that there is no significant difference in the bacterial community structure between locations from Eastern Tyrol and South Tyrol. The hierarchical clustering showed the segregation among the different Dolomites regions which was strongly pronounced in the case of VT populations when compared to ET and ST populations (Figure S4).

### No difference between the microbial communities of epidemic and endemic populations

To investigate potential differences in the bacterial community composition between populations in different epidemiological phases we compared the microbial communities between endemic (ET2 and ET4) and epidemic populations (ET1 and ET3) from Eastern Tyrol on a small local scale. The bacterial community composition was similar between the epidemic and endemic populations at both phylum and genus levels (Figure 6A; 6B). The bacterial species richness and diversity were similar between groups (Welch’s t-test: Chao1, t = 0.51824, df= 39.199 p = 0.6072; Shannon, t = 0.24332, df = 41.578, p = 0.809). Moreover, Canonical Analysis of Principal coordinates based on unweighted Unifrac and Bray-Curtis dissimilarities showed a strong overlap between the endemic and epidemic populations (Figure 6C; 6D). This was confirmed by Adonis multivariate analysis of variance, showing that there was no significant difference between the bacterial communities of endemic and epidemic populations of *I. typographus* (PERMANOVA, df= 1, F= 0.4194, R2= 0.00904, p = 0.7363). These results might indicate that the bacterial community is rather influenced by the geographical region than the epidemic status. However, several taxa had different relative abundances between epidemic and endemic populations. In particular, *Pseudoxanthomonas*, *Spiroplasma* and *Brevimonas* were more abundant in epidemic populations, whereas *Allorhizobium−Neorhizobium* and *Erwinia* were more abundant in endemic populations (Figure 6B).

**Figure 6:**
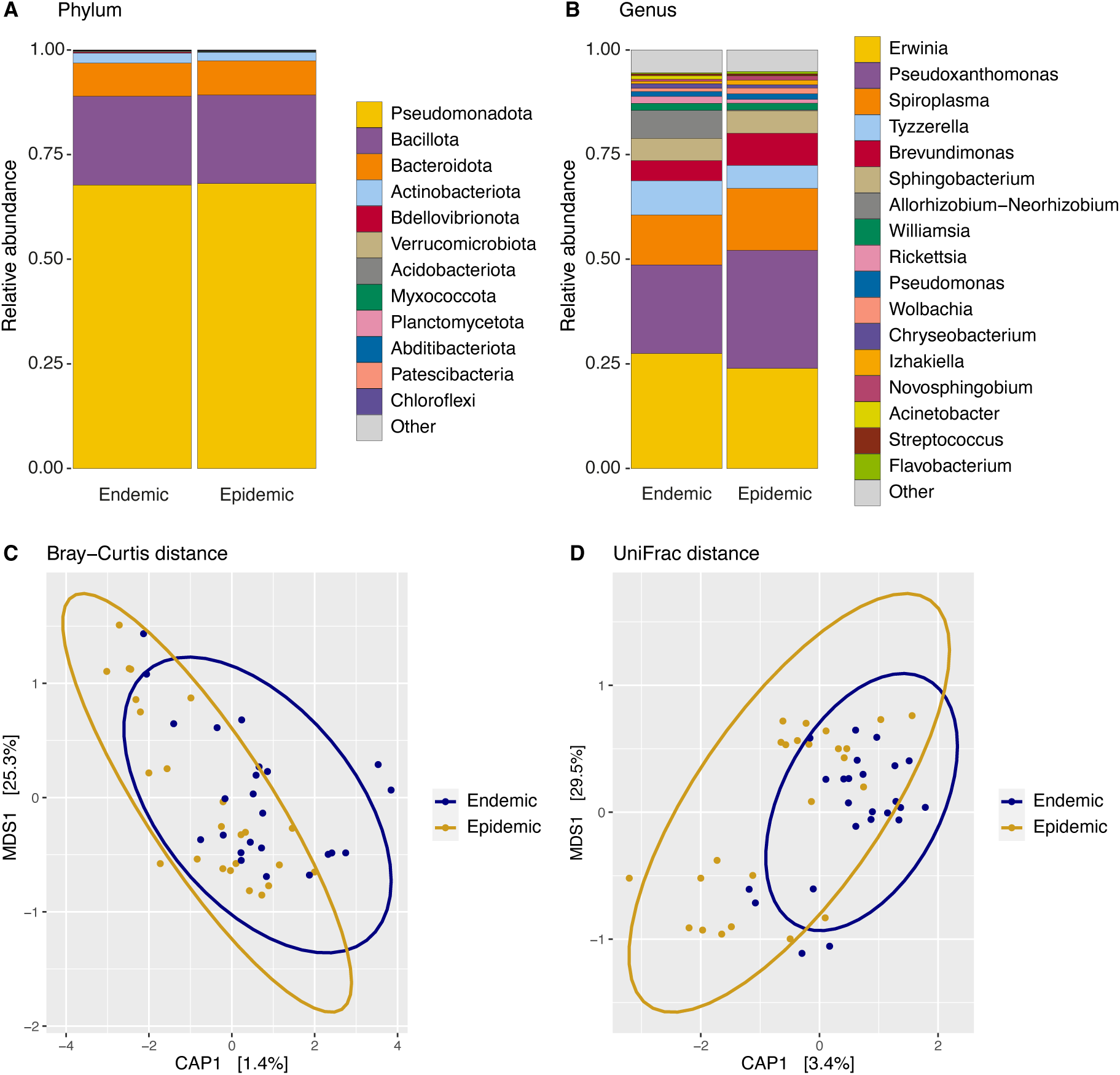
Bacterial community of endemic and epidemic populations of *I. typographus* from Eastern Tyrol at phylum level (A) and genus level (B). In (B) only the 17 most abundant genera were plotted. Canonical analysis of principal coordinates (CAP) based on Bray-Curtis dissimilarity (C) and unweighted UniFrac distance (D) between endemic and epidemic populations.

### Different bacterial communities between pre-overwintering and overwintering populations

To study the influence of overwintering on the bacterial community composition, we compared the microbial communities between pre-overwintering VT2, VT3 and VT6 (which were sampled in August and September 2020) and the overwintering VT4 and VT5 populations (which were sampled in December 2019 and January 2020). Despite sharing the same bacterial composition on the phylum level between pre-overwintering and overwintering individuals, there were differences in the abundance where *Allorhizobium-Neorhizobium* was more abundant in pre-overwintering and Erwinia, *Pseudomonas,* and *Pseudoxanthomonas* in overwintering populations. Particularly, the relative abundance of the genera *Acinetobacter, Serratia,* and *Spiroplasma* was significantly higher in pre-overwintering individuals. In contrast, the genera *Brevendomonas, Gordonia*, *Pajaroellobacter*, *Pedobacter,* and *Williamsia* were significantly more abundant in overwintering individuals (Figure S5). Both permutated multivariate ANOVA (Adonis) and pairwise comparison of PERMANOVA revealed a strong segregation between the pre-overwintering and overwintering individuals (all p < 0.001; Figure 7A; 7B).

**Figure 7:**
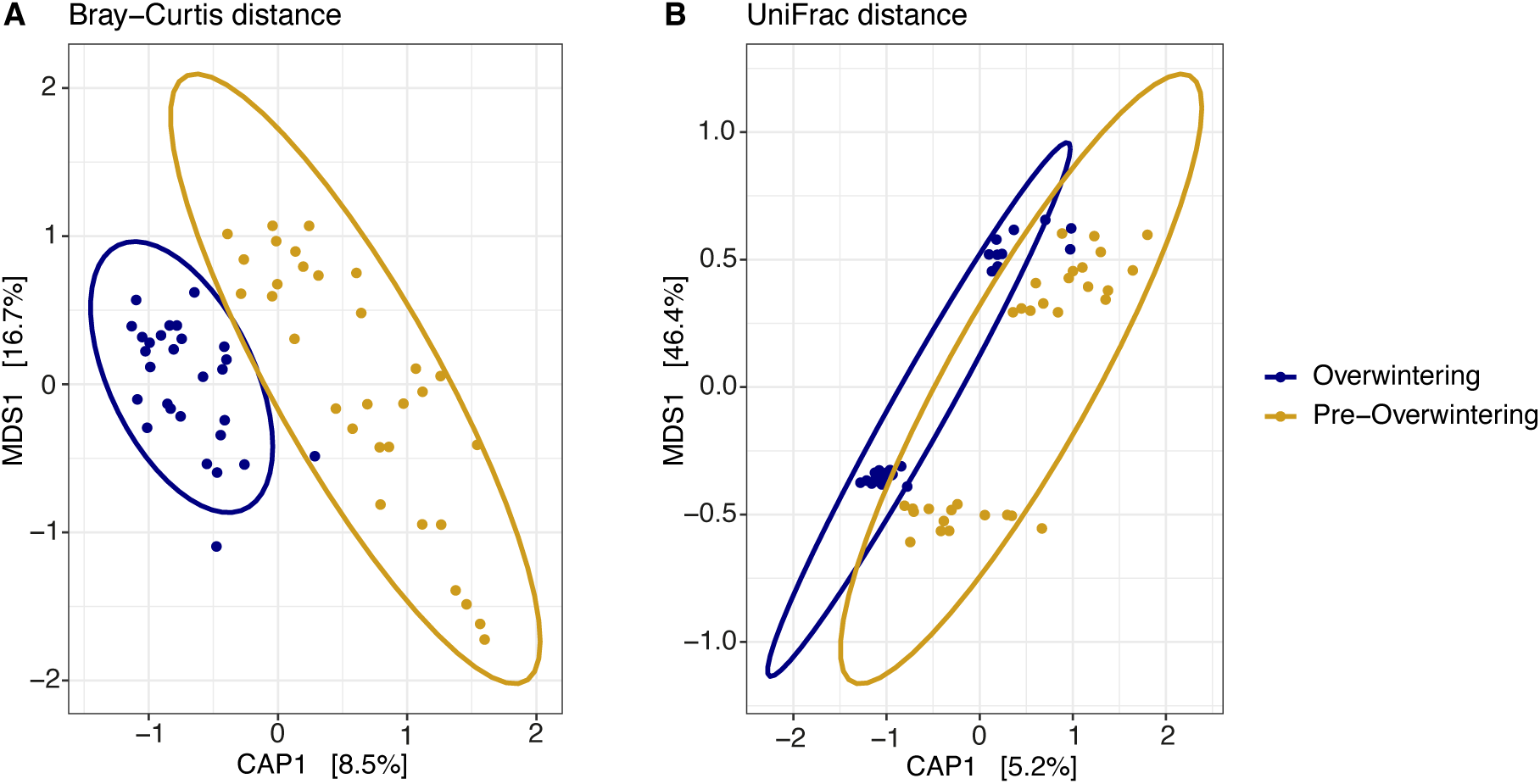
Canonical analysis of principal coordinates (CAP) based on Bray-Curtis dissimilarity (C) and UniFrac distance (D) between pre-overwintering and overwintering populations of *I. typographus* from Veneto.

### Predicted pathways of bacterial communities in I. typographus

Metagenome function predictions highlight differences among samples from different populations, regions, and season. The PICRUSt2 analysis predicted 419 gut microbial MetaCyc functional pathways showing a pattern across the different populations of *I. typographus* (Figure S6). Statistical analysis revealed that 333 (t-test/one-way ANOVA, FDR q < 0.05) and 59 biochemical pathways were differentially abundant across the different populations and between individuals of different overwintering phases, respectively. However, there was no statistical difference between the epidemiological phases. There were 26 biochemical pathways significantly enriched within the top 50 pathways divided by regions, 24 by populations, and 6 by season (Figure S6; Table S6). These included pathways related to the biosynthesis and degradation (of lipids, amino acids biosynthesis, as well as pathways mainly related to the “generation of precursor metabolites and energy”, and “nucleoside and nucleotide biosynthesis.”

The heatmap with the 44 most abundant varying pathways of this dataset is displayed, consistently showing patterns of abundance differences between the regions (Figure S6). The cluster A contains mainly pathways related to the synthesis of amino acids, which are generally more abundant among the samples of Eastern Tyrol, Russia (AI), Veneto (AII), while they are less abundant in South Tyrol (AII). The lipidic pathways enclosed in cluster B, was more abundant in the samples from Veneto. Conversely, they are less abundant in some populations from South Tyrol and Eastern Tyrol. Lastly, most of the pathways of cluster C related to energy and nucleotide biosynthesis are more common in all the regions except for Veneto. Within Veneto populations, biosynthesis of isoleucine (I, II, III), of valine, of branched amino acids, and pyruvate fermentation to isobutanol (engineered) significantly increased during overwintering (FDR q < 0.05) Moreover, fatty acids and beta-oxidation were also more common (raw-p < 0.05). Conversely, biosynthesis of gondoate, (5Z)-dodec-5-enoate, oleate IV, CDP-diacylglycerol (I, II), phospholipids, (Z)-octadec-11-enoate, palmitoleate, and fatty acid elongation-saturated significantly decreased (raw-p < 0.05). Additionally, in Eastern Tyrol lysine, threonine, and isoleucine I biosynthesis, was more present than in the other regions.

## Discussion

Here, we present a comprehensive assessment of different factors influencing the bacterial community composition of the European spruce bark beetle *I. typographus* across a wide geographic range including the four main postglacial refugial areas. Moreover, we specifically investigated possible changes of the bacterial community among populations in different epidemiological phases and beetles before and during overwintering in an area with a current bark beetle mass outbreak within the Dolomites.

While previous studies were based on pooled *I. typographus* guts from one restricted geographic area (Chakraborty et al. 2020; Fang et al. 2020; Yu et al. 2022; Veselská et al. 2023), our approach allowed a fine-scale analysis of several European populations on a single individual level. Our analysis revealed that the genera *Erwinia* and *Pseudoxanthomonas* which were previously described in *I. typographus* in the Czech Republic and in China (Chakraborty et al. 2020; Fang et al. 2020; Yu et al. 2022) can be considered the core bacterial community being present in all individuals (*Erwinia*) and in 98.44% of the individuals (*Pseudoxanthomonas*). The omnipresence of these two genera across most individuals from different populations across Europe suggests that these bacteria might be essential for *I. typographus* to successfully colonize trees and utilize the subcortical material as a food source. Since various other bark beetle species of the genera *Ips* (Chakraborty et al. 2020) and *Dendroctonus* (Adams et al. 2013) also harbored these bacteria, they might play an important role in the life histories of these widespread conifer bark beetles. Strains of *E. typographi* isolated from *I. typographus* were found to be tolerant to high concentrations of the monoterpene myrcene, one of the defensive compounds of Norway spruce (Skrodenytė-Arbačiauskienė et al. 2012), whereas members of the genus *Pseudoxanthomonas* contribute to cellulose and lignocellulose degradation and therefore play a potential role in the breakdown of plant cells in conifers (Kumar et al. 2015).

Apart from these two prevalent genera, other bacterial taxa were present in multiple populations, but at different frequencies. Since not all individuals harbor them, they might not be essential for the survival of the host but might play an additional important role for the beetle. For example, *Brevundimonas* and *Pseudomonas,* have been shown to contribute to the detoxification of conifer phytochemicals in *Dendroctonus* (Boone et al. 2013). While *Brevundimonas* is able to reduce diterpene abietic acid levels at low concentrations, two *Pseudomonas* species were shown to reduce concentrations of monoterpenes under controlled conditions (Boone et al. 2013). Since both taxa were present in all investigated populations, we assume that these bacterial taxa might be involved in overcoming host defenses and might therefore be especially important in colonizing healthy trees. Moreover, the genomic characterization of *Pseudomonas* strains isolated from *I. typographus* revealed various important pathways involved in the inhibition of entomopathogenic fungi as well as pathways for the hydrolyzation of cellulose, xylan, starch, and pectin (Peral-Aranega et al. 2020). Additionally, other dominant bacteria common among *Ips* species have been shown to be important for the biology of bark beetles: *Sphingobacterium* contributes to the decomposition of hemicellulose (Zhou et al. 2009) whereas *Acinetobacter* and *Williamsia* to the decomposition of lignin (Bugg et al. 2011). While *Sphingobacterium* and *Williamsia* were present in most individuals from all different regions, *Acinetobacter* was present in lower abundance in fewer individuals from most populations.

Glacial refugial areas during the Pleistocene have been shown to be important for the current geographic range and genetic structure of many species (Schmitt 2007). While most studies focused on how the glacial isolation of different populations shaped their population structure of different bark beetles (Bertheau et al. 2013), the potential effect on the microbial community has not been investigated. Moreover, a growing body of research supports the idea that microorganisms exhibit biogeographic patterns (Martiny et al. 2006) and therefore symbionts could be used to reconstruct the evolutionary history of their hosts. Here, we compared the bacterial communities of populations from the main putative refugial areas of *I. typographus* from the Apennines (Abetone), the Dinaric Alps (Croatia), the Carpathians (Rumania) and the Russian Plain (Stauffer et al. 1999; Bertheau et al. 2013), and observed significantly different bacterial communities between these regions. While the populations from the Apennines and Dinaric Alps clustered with the populations from Eastern Tyrol and South Tyrol, the populations from the Carpathian Mountains clustered with the populations from Veneto and Russia forming a unique cluster. Although we cannot exclude that ecological differences between the sampling years and sampling seasons had an impact on the outcome, our results indicate a potential influence of the Carpathian refugial area onto the Southern Alps, which is different from the genetic structure of *I. typographus* (Stauffer et al. 1999, Bertheau et al. 2013). Therefore, our study sheds new light on the evolution and post-glacial history of this bark beetle and its associated bacteria.

Although we found significant differences between the bacterial communities from different regions, we did not detect major differences between the bacterial communities of populations in different epidemiological phases, neither in terms of taxonomic richness and diversity nor community structure. These results suggest that *I. typographus* outbreaks might not be linked to a major shift in the bacterial community, at least not in the early phase of an outbreak. However, minor changes were observed for specific genera among the most abundant bacteria, such as *Pseudoxanthomonas* and *Spiroplasma* which had a higher relative abundance in the epidemic than in the endemic populations. While *Pseudoxanthomonas* was reported to have a role in the biodegradation of celluloses and lignocelluloses (Kumar et al. 2015) and might therefore play a role in digestion of xylem and phloem in the host (Fang et al. 2020), *Spiroplasma* can have various effects on their hosts, ranging from beneficial (such as increasing tolerance to natural enemies in their host to reproductive parasitism, particularly male-killing (Ballinger & Perlman 2019). It is unclear if and to which extant they influence the epidemiology of *I. typographus*.

An additional important factor in the biology of *I. typographus* is its overwintering behavior. *I. typographus* increases its cold tolerance and enters a reproductive diapause at the end of the season to increase the chance of overwinter survival (Annila 1969; Schebeck et al. 2017). Seasonal variations and diapause behavior might influence the bacterial community structure and the latter could support the insect in overcoming the harsh winter conditions, for instance by nutrient storage to build up energy reserves (Dittmer & Brucker 2021). A change in the bacterial community structure across seasons and during overwintering was observed in *Dendroctonus* bark beetles (Wang et al. 2017). By comparing populations sampled before and during the overwintering phase, we found a significant difference of the bacterial community. In particular, the genera *Acinetobacter*, *Seratia, Spiroplasma* and *Tyzzerella* were found at higher abundances in pre-overwintering beetles, while *Williamsia*, *Pseudomonas* and *Sphingobacterium* were more abundant in overwintering populations. The last two genera were also enriched during winter in the gut microbiota of *Dendroctonus armandi* larvae (Wang et al. 2017), highlighting a potential contribution to overwintering of bark beetles. It has been hypothesized that bacteria from the family *Sphingobacteriaceae* are associated with insect overwintering and survival at low temperatures (Wang et al. 2017), whereas *Pseudomonas* might be involved in increasing *I. typographus* resistance to fungal pathogens and thus increasing the chance of survival (Peral-Aranega et al. 2020). Additionally, the genus *Novosphingobium* was significantly more abundant in overwintering beetles. Cheng et al. (2018) reported the involvement of *Novosphingobium* isolates in naringenin degradation activity in red turpentine beetle.

To predict the functional capabilities of the bacterial community we performed a PICRUSt analysis and found several amino acids such as branched-chain-, aromatic-, isoleucine, lysine, valine, threonine, glycine, and serine that were predicted within the major biochemical pathways, likely supporting the fitness of *I. typographus* (García-Fraile, 2018). Isoleucine is an amino acid that is crucially needed to emit several aliphatic/aromatic alcohols and esters in fungi, including those associated with bark beetles (Lehenberger, 2021). Thus, its importance is ultimately linked to the chemical communication between microbe and bark beetle interactions. Beta oxidation reactions are involved in detoxification mechanisms of pine terpenes by bark beetle-associated fungi (Hammerbacher et al., 2019). Interestingly, bacteria seem to provide several lipid sources, which in the populations in Veneto are more abundant in pre-ovewintering individuals. These could be then used by the beetle as fat reserve for the following months and in preparation for overwintering (Botterweg 1983). Lastly, our data show several similarities with the recent findings on the predicted bacterial pathways of the *I. typographus* pathosystem (Chakraborty et al., 2023). This implies that such pathways not only can be common to adult beetles from different regions, but also to the larvae and galleries, suggesting conserved functions of the bacteriome associated with this pest.

In conclusion, our study on the bacterial community of *I. typographus* across different geographic regions and epidemiological and seasonal phases highlights the important role of bacteria in this pest species. Especially the presence of *Erwinia* and *Pseudoxanthomonas* in all analyzed individuals suggests their essential role, likely as obligatory symbionts for this bark beetle. Further studies are needed to investigate their influence on the host and transmission route to the offspring. Similarly, the role of several less-frequent genera associated with *I. typographus* need to be further investigated to understand their role for its host.

## Acknowledgements

We thank A. Andriolo, H. Lercher, A. Stradner and F. Simões for technical and analytical supports, PHW. Biedermann, J. Carlos Cambronero Heinrichs for scientific advises and the editors Antonio Biondi and Nicolas Desneux as well as two anonymous reviewers for helpful comments on the manuscript. Financial support was provided by INTERREG Dolomiti Live (projects ITAT4132 and ITAT4153), the province of Bolzano, the Hermann Rubner Privatstiftung Onlus, and the Amt der Tiroler Landesregierung Abteilung Waldschutz for funding supports. We thank the Department of Innovation, Research and University of the Autonomous Province of Bozen/Bolzano for covering the open access publication charges.

