## Supplementary figures and images for "The bacterial community of the European spruce bark beetle in space and time"

### Figure S1

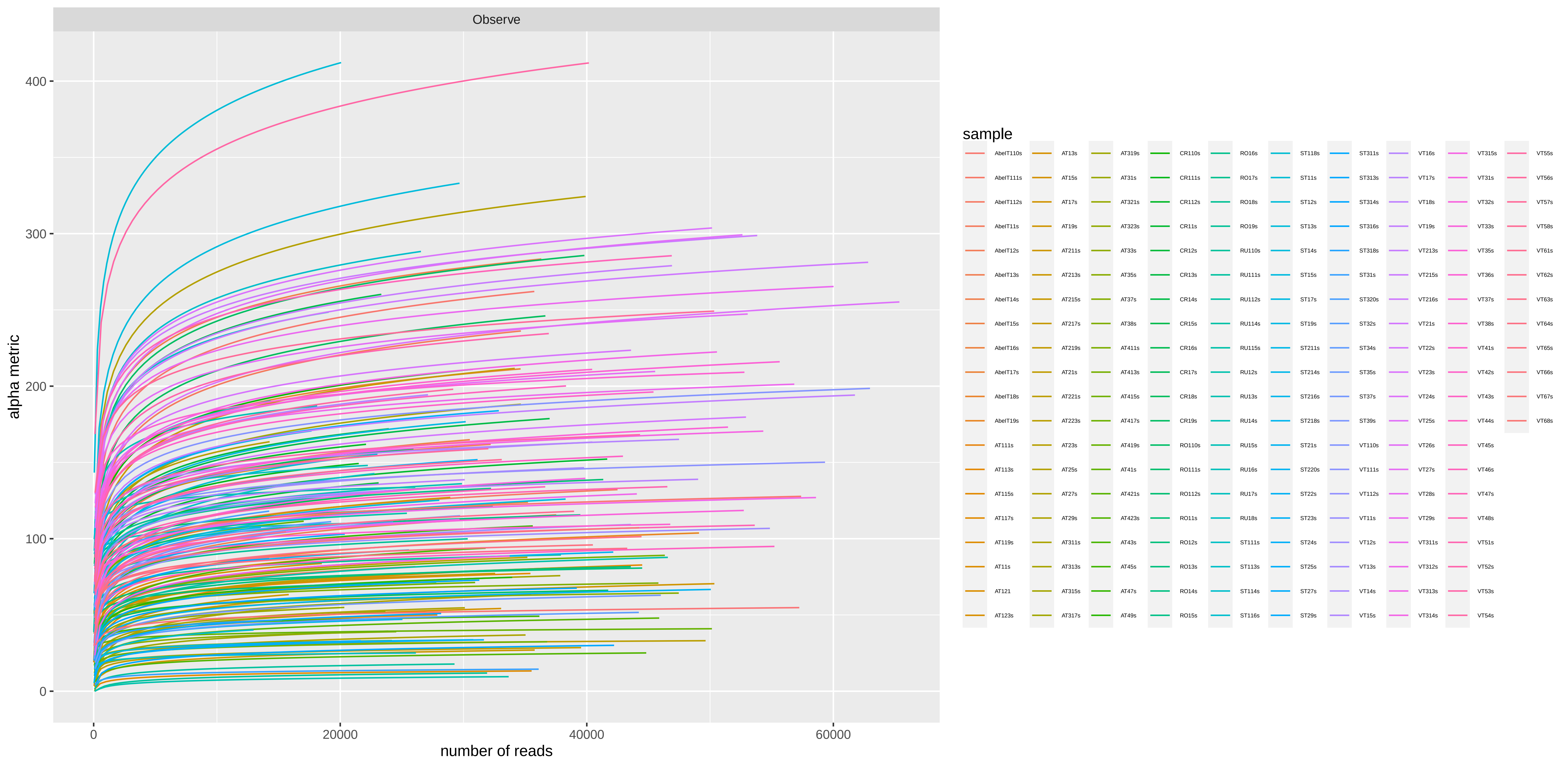

### Figure S2

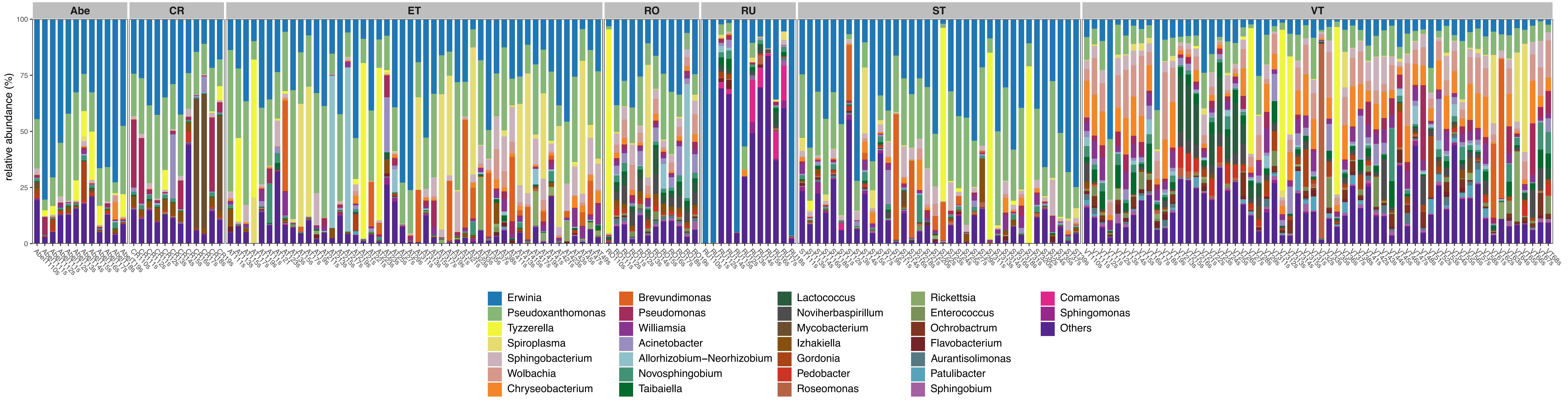

### Figure S3

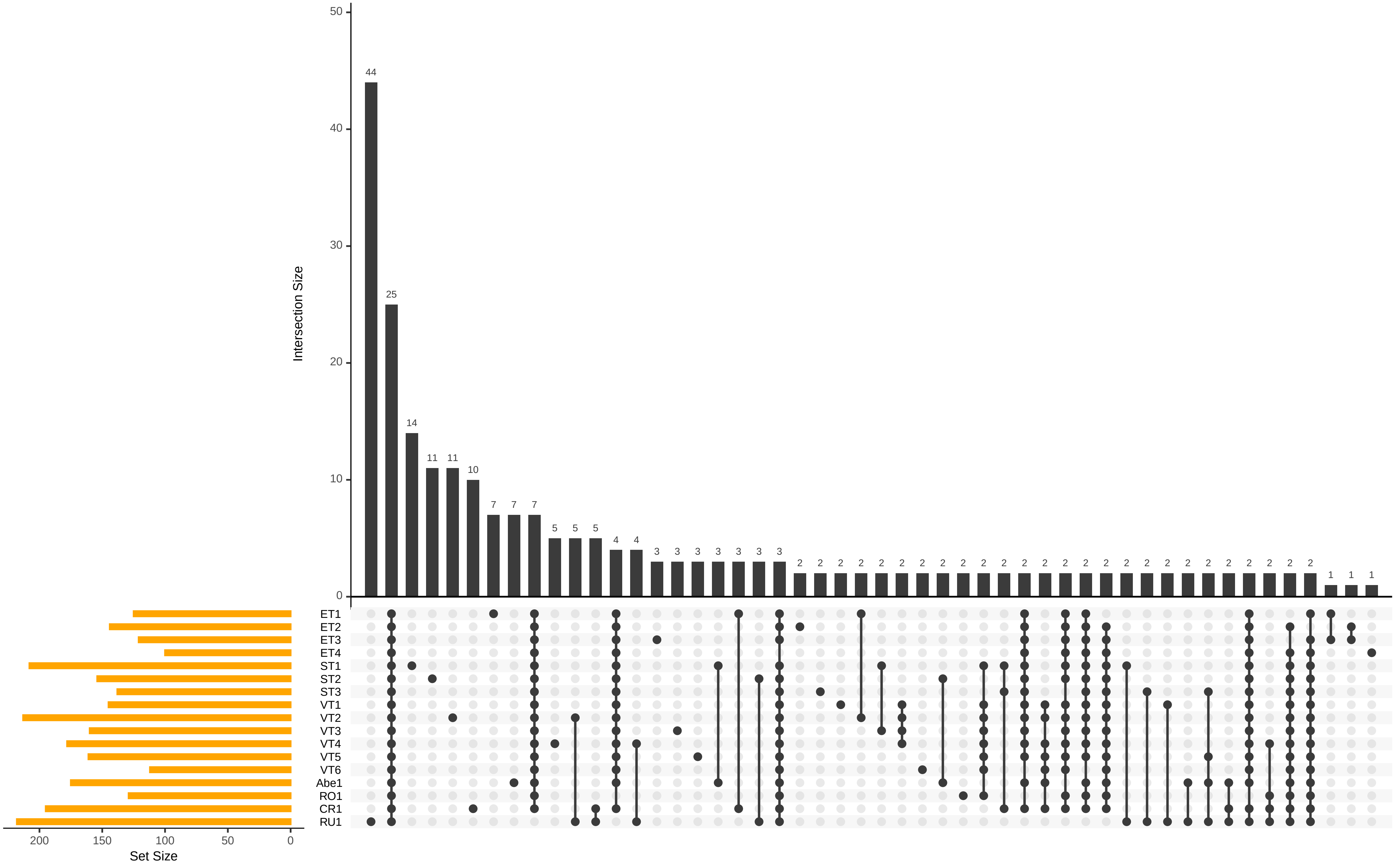

### Figure S4

#### Circular Hierarchical Cluster (UniFrac)

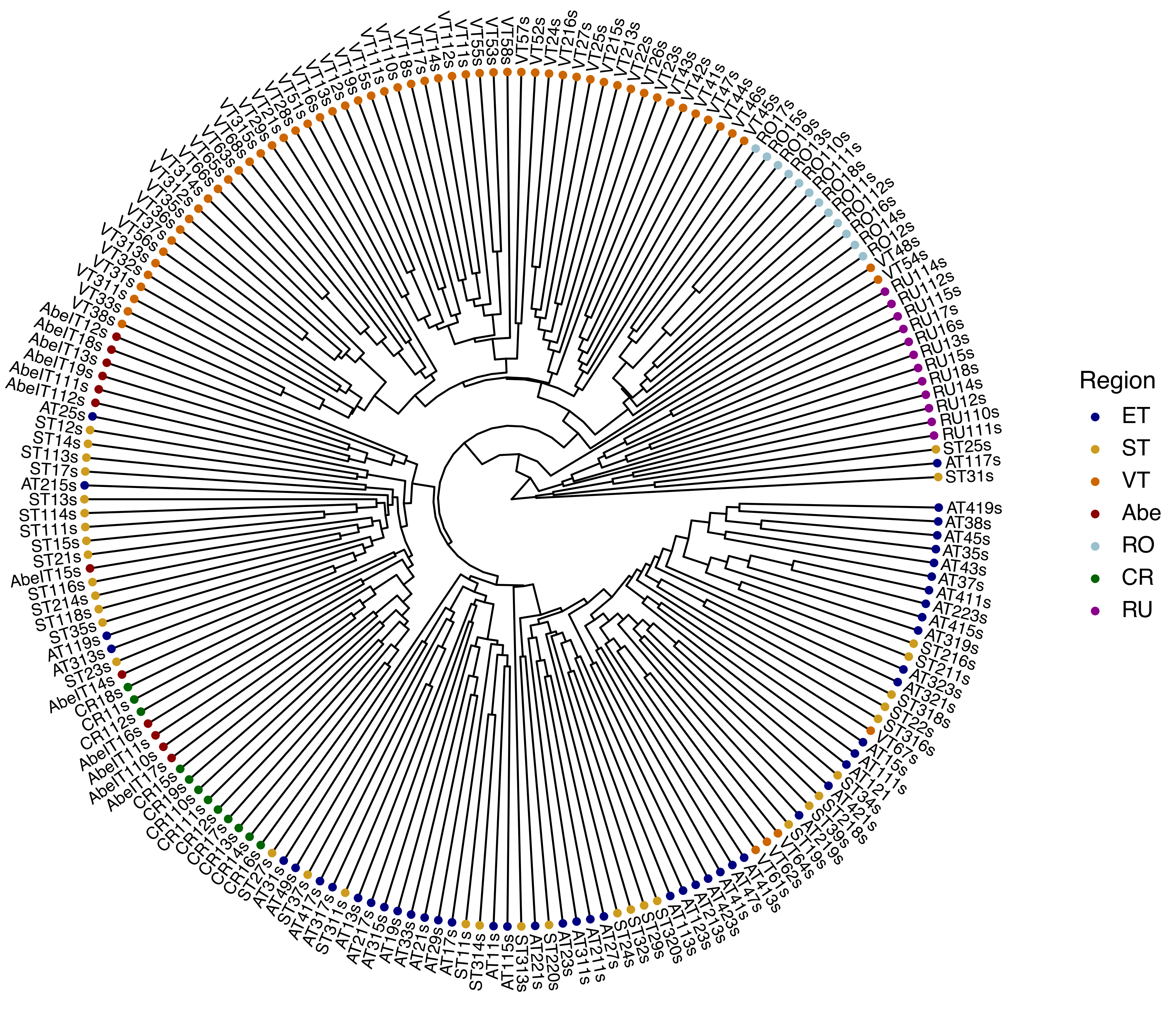

### Figure S5

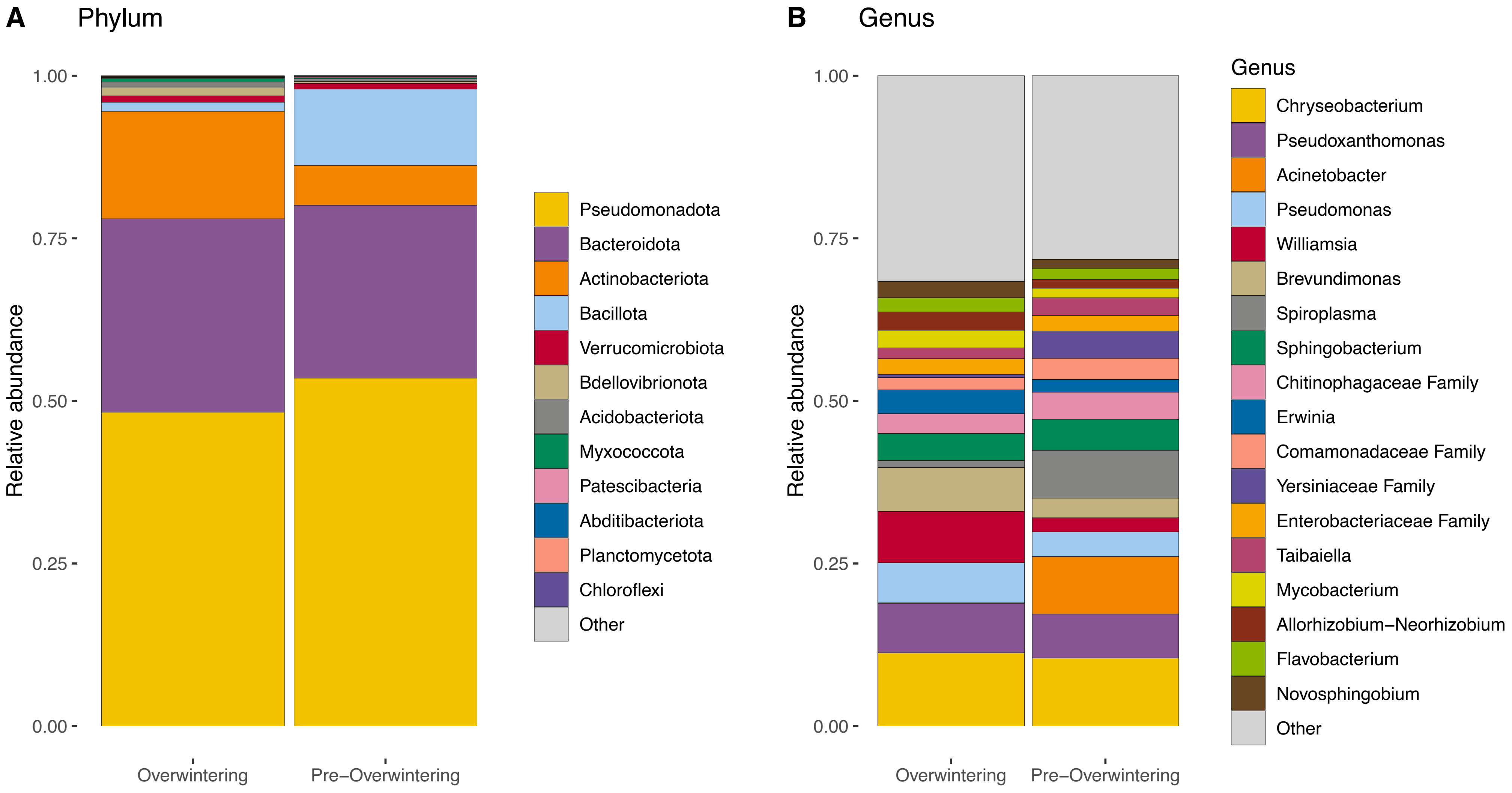

### Figure S6

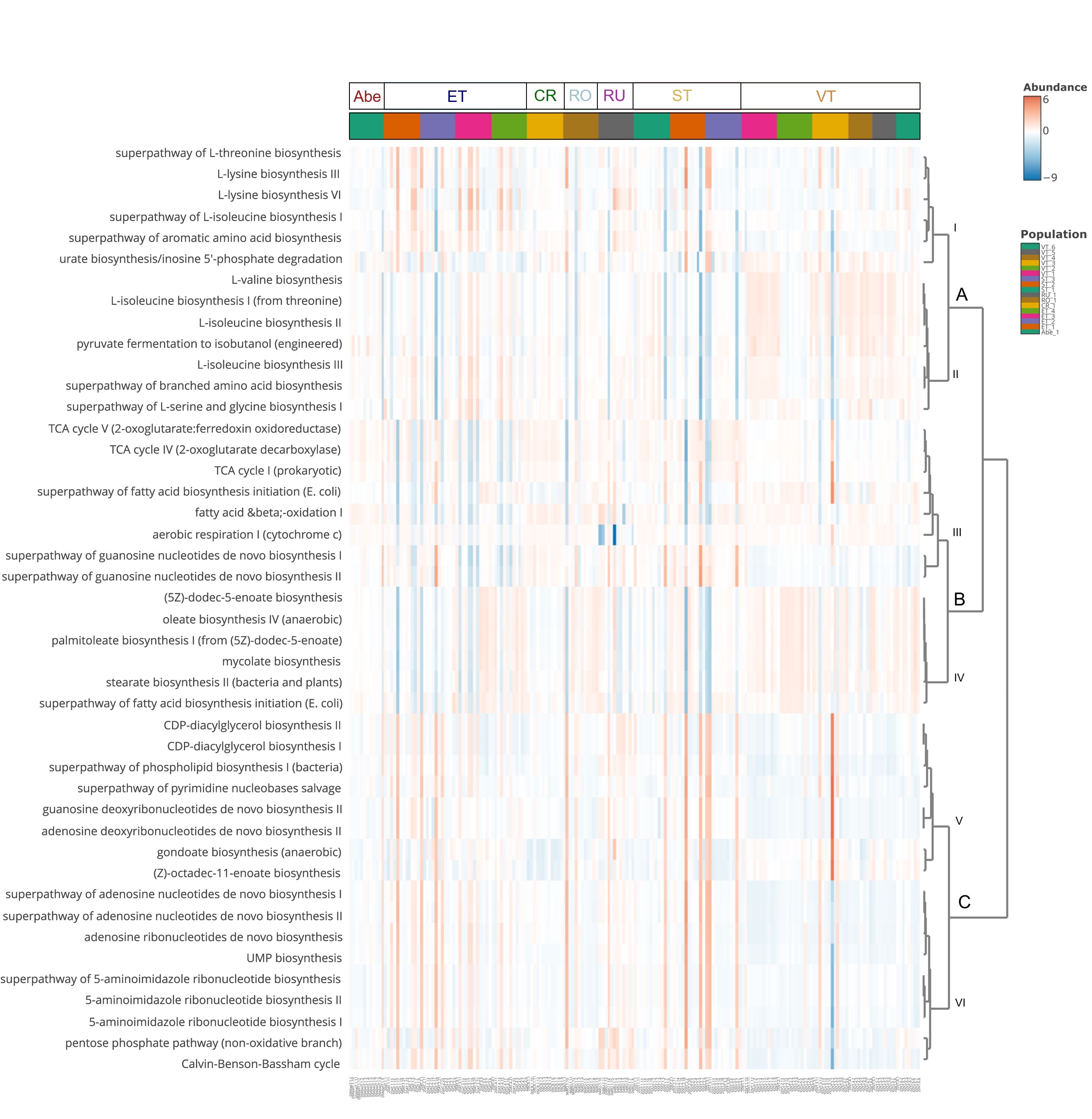
