## Supplementary material for "The bacterial community of the European spruce bark beetle in space and time": Table S1

**Table S1:** List of all sampling locations including the population code, latitude/longitude, number of analyzed individuals, epidemiological and overwintering status of *Ips typographus* and date of collection (month, year).

| Locality | Population | Latitude/Longitude | no | Status | month | Year |
| --- | --- | --- | --- | --- | --- | --- |
| Eastern Tyrol (Austria) | ET1 (Kals) | 46°56'54.34"N  12°36'15.35"E | 12 | Epidemic | July | 2020 |
|  | ET2 (Kals) | 46°57'28.78"N  12°37'12.69"E | 12 | Endemic | July | 2020 |
|  | ET3 (Obertilliach) | 46°41'59.26"N  12°36'0.32"E | 12 | Epidemic | July | 2020 |
|  | ET4 (Obertilliach) | 46°43'59.93"N  12°33'0.11"E | 12 | Endemic | July | 2020 |
| South Tyrol (Italy) | ST1 (San Candido) | 46°44'34.42"N  12°20'11.85"E | 12 |  | July | 2020 |
|  | ST2 (San Candido) | 46°44'35.26"N  12°22'4.43"E | 12 |  | July | 2020 |
|  | ST3 (Riva di Tures) | 46°53'17.41"N  11°52'21.19"E | 12 |  | July | 2020 |
| Veneto (Italy) | VT1 (Paluzza) | 46°31'41.97"N  13° 1'15.73"E | 12 |  | February | 2020 |
|  | VT2 (Col di Prà) | 46°17'45.90"N  11°55'33.63"E | 12 | Pre-Overwintering | August | 2020 |
|  | VT3 (Vigo di Cadore) | 46°30'28.66"N  12°29'1.33"E | 12 | Pre-Overwintering | September | 2020 |
|  | VT4 (Forni di sopra) | 46°19'59.88"N  12°19'30.72"E | 8 | Overwintering | January | 2020 |
|  | VT5 (Giazza) | 45°39'15.44"N  11° 7'30.98"E | 8 | Overwintering | December | 2019 |
|  | VT6 (Selva di Progno) | 45°40'41.47"N  11° 6'59.92"E | 8 | Pre-Overwintering | September | 2020 |
| Abetone (Italy) | Abe1 | 44° 8'0.00"N  10°38'60.00"E | 12 |  |  | 2009 |
| Romania | RO1 (Belis) | 46° 3'54.83"N  23° 0'46.60"E | 12 |  |  | 1996 |
| Croatia | CR1 (Vrhovine) | 44°50'48.82"N  15°25'24.28"E | 12 |  |  | 2009 |
| Russia | RU1 (Alexandrow) | 56°22'34.82"N  38°47'19.26"E | 12 |  |  | 1995 |
