## Supplementary material for "The bacterial community of the European spruce bark beetle in space and time": Table S3

Table S3. Pairwise comparison of Chao1 index of *I. typographus* collected from different localities.

| Region | diff | lwr | upr | p adj |
| --- | --- | --- | --- | --- |
| CR-Abe | 10 | -40.972875 | 60.972875 | 0.9971819 |
| ET-Abe | -57.0625 | -97.360096 | -16.764904 | 0.0007429 |
| RO-Abe | -50.416667 | -101.38954 | 0.5562084 | 0.0546804 |
| RU-Abe | -68.5 | -119.47288 | -17.527125 | 0.0017065 |
| ST-Abe | -30.722222 | -72.3414 | 10.896956 | 0.3002883 |
| VT-Abe | -6.966667 | -46.450086 | 32.5167525 | 0.9984439 |
| ET-CR | -67.0625 | -107.3601 | -26.764904 | 0.0000323 |
| RO-CR | -60.416667 | -111.38954 | -9.4437916 | 0.0092023 |
| RU-CR | -78.5 | -129.47288 | -27.527125 | 0.0001632 |
| ST-CR | -40.722222 | -82.3414 | 0.896956 | 0.0595967 |
| VT-CR | -16.966667 | -56.450086 | 22.5167525 | 0.8598785 |
| RO-ET | 6.645833 | -33.651763 | 46.9434293 | 0.9989386 |
| RU-ET | -11.4375 | -51.735096 | 28.860096 | 0.9796354 |
| ST-ET | 26.340278 | -1.188221 | 53.8687766 | 0.0707335 |
| VT-ET | 50.095833 | 25.917276 | 74.2743909 | 0.0000001 |
| RU-RO | -18.083333 | -69.056208 | 32.8895417 | 0.9394407 |
| ST-RO | 19.694444 | -21.924734 | 61.3136226 | 0.7955452 |
| VT-RO | 43.45 | 3.966581 | 82.9334192 | 0.0207721 |
| ST-RU | 37.777778 | -3.8414 | 79.396956 | 0.1025407 |
| VT-RU | 61.533333 | 22.049914 | 101.016753 | 0.0001291 |
| VT-ST | 23.755556 | -2.566724 | 50.077835 | 0.1064998 |
