## Supplementary material for "The bacterial community of the European spruce bark beetle in space and time": Table S4

Table S4. Pairwise comparison of Shannon index of *I. typographus* collected from different localities.

| Region | diff | lwr | upr | p adj |
| --- | --- | --- | --- | --- |
| CR-Abe | 0.38657681 | -0.4761482 | 1.2493019 | 0.834226 |
| ET-Abe | -0.0751452 | -0.7571892 | 0.6068989 | 0.999897 |
| RO-Abe | 0.67544026 | -0.1872848 | 1.5381653 | 0.2336237 |
| RU-Abe | 0.309467 | -0.5532581 | 1.1721921 | 0.936262 |
| ST-Abe | 0.08668196 | -0.6177301 | 0.791094 | 0.9998035 |
| VT-Abe | 1.19724651 | 0.52898256 | 1.8655105 | 0.0000056 |
| ET-CR | -0.461722 | -1.143766 | 0.220322 | 0.4068381 |
| RO-CR | 0.28886345 | -0.5738616 | 1.1515885 | 0.9538804 |
| RU-CR | -0.0771098 | -0.9398349 | 0.7856152 | 0.9999699 |
| ST-CR | -0.2998949 | -1.0043069 | 0.4045172 | 0.8651239 |
| VT-CR | 0.81066969 | 0.14240574 | 1.4789336 | 0.0069494 |
| RO-ET | 0.75058544 | 0.06854141 | 1.4326295 | 0.0207659 |
| RU-ET | 0.38461219 | -0.2974319 | 1.0666562 | 0.6295368 |
| ST-ET | 0.16182715 | -0.3040976 | 0.6277519 | 0.9451423 |
| VT-ET | 1.27239169 | 0.86316527 | 1.6816181 | 0.0001 |
| RU-RO | -0.3659733 | -1.2286983 | 0.4967518 | 0.8671134 |
| ST-RO | -0.5887583 | -1.2931704 | 0.1156538 | 0.168387 |
| VT-RO | 0.52180625 | -0.1464577 | 1.1900702 | 0.236487 |
| ST-RU | -0.222785 | -0.9271971 | 0.481627 | 0.9649998 |
| VT-RU | 0.8877795 | 0.21951555 | 1.5560435 | 0.0020268 |
| VT-ST | 1.11056455 | 0.66505525 | 1.5560738 | 0.0001 |
