## Supplementary material for "The bacterial community of the European spruce bark beetle in space and time": Table S5

Table S5. Pairwise comparison of *I. typographus* bacterial community structure collected from different localities based on unweighted UniFrac distance matrix at 999 permutations.

| Locality | Sum Of Sqs | R2 | F | Pr(>F) | sig |
| --- | --- | --- | --- | --- | --- |
| RU-Abe | 0.11434 | 0.23569 | 6.784 | 0.000999 | ** |
| RU-CR | 0.50937 | 0.24151 | 7.0052 | 0.000999 | *** |
| RU-ET | 0.4032 | 0.10586 | 6.8667 | 0.001998 | ** |
| RU-ST | 0.25399 | 0.10518 | 5.4069 | 0.007992 | ** |
| RU-VT | 0.5330 | 0.16822 | 14.157 | 0.000999 | *** |
| RU-RO | 0.15780 | 0.23358 | 6.7048 | 0.000999 | *** |
| Abe-CR | 0.36085 | 0.20156 | 5.5538 | 0.000999 | *** |
| Abe-ET | 0.2752 | 0.07839 | 4.9337 | 0.006993 | ** |
| Abe-ST | 0.1058 | 0.05047 | 2.445 | 0.05694 | . |
| Abe-VT | 0.55536 | 0.18385 | 15.768 | 0.000999 | *** |
| Abe-RO | 0.16637 | 0.32378 | 10.534 | 0.000999 | *** |
| CR-ET | 0.2854 | 0.06009 | 3.7083 | 0.02198 | * |
| CR-ST | 0.2846 | 0.08122 | 4.0665 | 0.02198 | * |
| CR-VT | 0.9598 | 0.20624 | 18.187 | 0.000999 | *** |
| CR-RO | 0.45706 | 0.22477 | 6.3788 | 0.000999 | *** |
| ET-ST | 0.1129 | 0.02197 | 1.8422 | 0.1449 |  |
| ET-VT | 1.2587 | 0.18623 | 24.257 | 0.000999 | *** |
| ET-RO | 0.2883 | 0.07853 | 4.9431 | 0.01299 | * |
| ST-VT | 1.0064 | 0.19127 | 22.232 | 0.000999 | *** |
| ST-RO | 0.21708 | 0.0922 | 4.6717 | 0.01199 | * |
| VT-RO | 0.14357 | 0.05209 | 3.847 | 0.01099 | * |

Significance levels .0 ‘***’, 0.001 ‘**’, 0.01 ‘*’, 0.05 ‘.’
